# A computational framework for converting high-throughput DNA sequencing data into neural circuit connectivity

**DOI:** 10.1101/244079

**Authors:** Hassana Oyibo, Cang Cao, Daniel D. Ferrante, Huiqing Zhan, Alex Koulakov, Lynn Enquist, Joshua Dubnau, Anthony Zador

## Abstract

There is growing interest in determining the connectivity of neural circuits—the “connectome”—at single neuron resolution. Most approaches to circuit mapping rely on either microscopy or physiology, but these approaches have very limited throughput. We have recently proposed BOINC (Barcoding of Individual Neuronal Connectivity), a radically different approach to connectivity mapping based on high-throughput DNA sequencing. Here we describe the set of computational algorithms that serve to convert sequencing data into neural connectivity. We apply our computational pipeline to the results of proof-of-principle experiments illustrating an implementation of BOINC based on pseudorabies virus (PRV). PRV is capable of traversing individual synapses and carry DNA barcodes from one cell to another. Using this high-throughput sequencing data, we obtain 456-by-486 connectivity matrix between putative neurons. An inexpensive high-throughput technique for establishing circuit connectivity at single neuron resolution would represent a major advance in neuroscience.

## Introduction

The mouse brain consists of tens of millions of neurons each connected by thousands of synapses; the human brain has tens of thousands of times more. The details of these connections are crucial in determining brain function. Malformation of these connections during prenatal and early postnatal development can lead to mental retardation, autism or schizophrenia (Dani and Nelson, 2009; Rinaldi et al., 2008; Testa-Silva et al., 2012); loss of specific connections later in life is associated with neurodegenerative diseases such as Alzheimer’s. An efficient method for determining the brain’s wiring diagram at single neuron resolution—its *connectome*—would represent a major advance in neuroscience.

Current approaches to determining the connectome are based on electron microscopy. So far, the complete connectome has been established for only one organism, the tiny worm *C. elegans,* with 302 neurons connected by about 7000 synapses. However, determining the connectome of even this simple nervous system was a heroic task, requiring over 50 person-years of labor. Although recent technological advances (Bock et al., 2011; Briggman et al., 2011) have led to considerable increases in throughput, electron microscopic approaches to reconstructing neural circuitry remain slow, expensive and laborious.

To circumvent the challenges associated with determining the connectome by microscopy, we have been pursuing an alternative strategy in which neural connectivity is converted into a form that can be decoded by high-throughput DNA sequencing (Zador et al., 2012). The appeal of sequencing is that it is fast and inexpensive—the cost of sequencing a human genome (~3B nucleotides) is approaching $1000. Thus by converting brain connectivity from a problem of microscopy to a problem of sequencing, we render it tractable using current techniques.

To convert connectivity into a sequencing problem, three challenges must be solved. *First,* each neuron must be endowed with a unique DNA or RNA “barcode.” The potential diversity of sequences of DNA grows exponentially with the sequence length, so that even very short sequences can be used to barcode neuronal populations uniquely. For example, a barcode consisting of a random string of 25 nucleotides has a potential diversity of 4^25^ = 10^15^, far more than the number of neurons (<10^8^ neurons) in a mouse brain. *Second,* barcodes from synaptically connected neurons must be associated into barcode pairs representing connections between neurons. *Third,* synaptically associated barcodes must be sequenced. The barcode pairs associated at synapses will then contain an imprint of synaptic connectivity of the entire network with single-cell resolution. This connectivity can be obtained by sequencing DNA barcodes using conventional next-generation sequencing technology, or by in situ sequencing methods such as BaristaSeq (Chen et al., 2017).

Here we present computational methods that can be used to reconstruct connectivity based on experimental data obtained in neuronal co-culture. The experimental approach adopted here is our first generation system. This system uses pseudorabies virus (PRV), a double stranded DNA virus of the herpes family, to move barcodes between connected neurons. PRV has evolved exquisite mechanisms for moving genetic material between functionally connected neurons, and has been used extensively for tracing neural circuits (Card and Enquist, 2012). Although there are technical limitations arising from our first generation transsynaptic approach which are mitigated in our second generation approach [synseq; (Peikon et al., 2017)], our results demonstrate the feasibility of each of the key components for converting neuronal connectivity into a form that can be read by next generation sequencing.

## Materials and Methods

### DNA constructs and Barcode library

PRV amplicons are plasmids containing a PRV origin of replication (ORI) and no other viral genes. In the presence of a PRV strain that can replicate, the amplicons replicates to high copy number. If the PRV amplicon also contains a viral DNA packaging sequence (a PAC site), the replicated amplicon will be packaged efficiently in helper virus infected cells into virus particles that can infect cells and deliver the amplicon DNA. We constructed two types of amplicons called *hosts* and *invaders.* The host amplicon did not contain a PRV PAC sequence and therefore could not be packaged into virions produced by the helper PRV. The invader replicon contained both an ORI and a PAC site. To construct the invader amplicon vector, PRV ORI and PAC sequences were amplified from pORI-PAC-GFP (a generous gift from Dr. Enrique Tabarés) and subcloned together with CAGGS-dsRed Express into pSMART vector. To construct the host amplicon vector, PRV ORI was amplified from pORI-PAC-GFP and subcloned together with CAGGS-GFP into the pSMART vector. The phiC31 plasmid pCS-P (http://www.ncbi.nlm.nih.gov/pmc/articles/PMC2834998/) was a generous gift from Dr. Michele P. Calos (Stanford). The oligonucleotides used for invader and invader barcode library construction were synthesized by IDT (Coralville, IA), and made double stranded by primer annealing and extension. The invader amplicons contained the Solexa II sequence from Illumina followed by 25 random nucleotides that are then followed by a library accession tag sequence (TAGC) and the truncated phiC31 integration sequence attB. The invader oligonucleotide sequence was as follows:

> CAAGCAGAAGACGGCATACGAGATCGGTCTCGGCATTCCTGCTGAACCGCTCTTCCGATCT-N25-TAGCGTGCGGGTGCCAGGGCGTGCCCTTGGGCTCCCCGGGCGCGTACTCC.

The host contained the Solexa I sequence from Illumina followed by 25 random nucleotides, followed by a library accession tag sequence (GTCA) and the truncated phiC31 integration attP sequence. The host oligonucleotide sequence was as follows:

> AATGATACGGCGACCACCGAGATCTACACTCTTTCCCTACACGACGCTCTTCCGATCT-N25-GTCAACGCCCCCAACTGAGAGAACTCAAGGGCACGCCCTGGCACCCGCAC.

EcoRI and NheI sites were present at the ends of both oligonucleotides, and the digested oligonucleotides were ligated into their respective amplicon vectors. The resulting plasmid library was then transformed into bacteria and amplified. Assuming that each bacterial colony contained a single unique barcode, the diversity of each library was estimated by counting the number of colonies obtained from a small aliquot of transformed bacteria, and in some cases validated by high-throughput sequencing. Library diversity was typically in the range of 10^6^–10^7^.

### Cell culture, electroporation and infection

Embryonic day 18 rat cortical tissue was purchased from Brainbits (Springfield, IL). After 20 minute-digestion in papain, the cortical tissue was resuspended in NbActiv4 medium (Brainbits, Springfield, IL.) and subjected to electroporation. A suspension of 5×10^5^ cells was electroporated with 1μg of the invader or host amplicon ± 1 μg of phiC31 plasmids using an Amaxa Nucleofector (Amaxa Biosystems, Lonza) following the manufacturer’s directions. After electroporation, cells with invader or host amplicons ± phiC31 were mixed and co-plated in a 24-well tissue culture plate pre-coated with poly-L-lysine (Sigma, St. Louis, MO). Three or 13 days after co-plating, cells were infected by the Bartha strain of PRV at a multiplicity of infection (MOI) of 1.

### *In utero* electroporation and infection

*In utero* electroporations were performed as described elsewhere (Dixit et al., 2011). Briefly: the amplicon plasmid electroporated included PRV-PAC, PRV-ORI and a GFP expression cassette. The amplicon DNA was prepared with a Qiagen endotoxin free maxi prep, and injected at a concentration of ~1μg/μl after mixing with ~0.025% fast green dye. Pulled glass capillaries with an opening of ~10 μm were used to deliver the DNA. Pregnant FVB mice 16 days into gestation were anesthetized with isoflurane, their uterus was exposed and approximately 1–2μls of the amplicon plasmid mix was microinjected using a Picospritzer II (Parker Instruments), into the lateral ventricles of the cerebral cortex through the uterus. Electroporation was achieved by unipolar discharge of 50V in five 50ms pulses spaced 950ms apart using a BTX ECM 830 Square Wave Electroporator. The positive charged paddle was placed on the left side of the pup brain to target amplicons plasmid to left cortex. After suturing, the pregnant mice were placed on heating pads until they awoke. At P23 electroporated pups were injected in the left auditory cortex with ~200nl of PRV Becker titer 10^9^, Brains were harvested following transcardial perfusion 24 hours after infection. All animal protocols were in accordance with the Cold Spring Harbor Laboratory Animal Care and Use Committee and carried out in accordance with National Institutes of Health standards.

### DNA extraction, PCR and sequencing

DNA was extracted using the Zymo viral DNA kit (Zymo research, Irvine, CA) according to the manufacturer’s instructions. The fused barcode pairs were amplified by PCR using Phusion Flash PCR Master Mix (Finnzymes, Vantaa, Finland) under the following conditions: 98°C for 10 sec, 37× [98°C for 1 sec, 72°C for 40 sec], 72°C for 1 min. 1 μl of the PCR product was subcloned into blunt TOPO vector (Invitrogen, Carlsbad, CA) and sequenced using conventional Sanger methods the remainder was purified and sequenced by Illumina (GA-II).

### Force directed graph drawing

To visualize the network in space, we minimized the following functional with respect to the 3D positions of network nodes 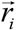 (Fruchterman and Reingold, 1991)

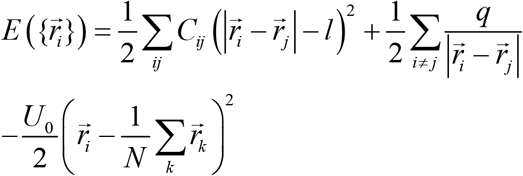

Here *N* is the total number of nodes (456 + 786 = 1242 in Figure 8, for example) *C_ij_* is the connectivity matrix between nodes *i* and *j* (1242×1242 for Figure 8), *l* = 1 is the equilibrium spring length, *q* = 0.016 is the repulsive constant, and *U*_0_ = 0.01. Three terms in this sum describe elastic springs between connected nodes, electrostatic repulsion between all nodes, and repulsion from the center of mass, respectively. We minimized the functional using the conjugate gradient algorithm (Saad, 2003). This yielded positions of the nodes in the minimum 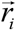 that are shown in Figures 8 and 11.

## Results

We developed methods to accomplish each of the three steps needed to convert neuronal connectivity into a DNA sequencing problem (Fig. 1). First, we developed a strategy to deliver DNA barcodes to neurons using PRV amplicons. Second, we devised a means to move amplicons. After amplicon movement to synaptically connected neurons, each neuron contains copies of its own barcoded amplicon, as well as copies of barcoded amplicons from synaptic partners. Finally, we used phiC31 integrase (Groth et al., 2000) to join the barcodes from synaptically connected partners. The joined barcode pairs are then sequenced.

**Figure 1.**
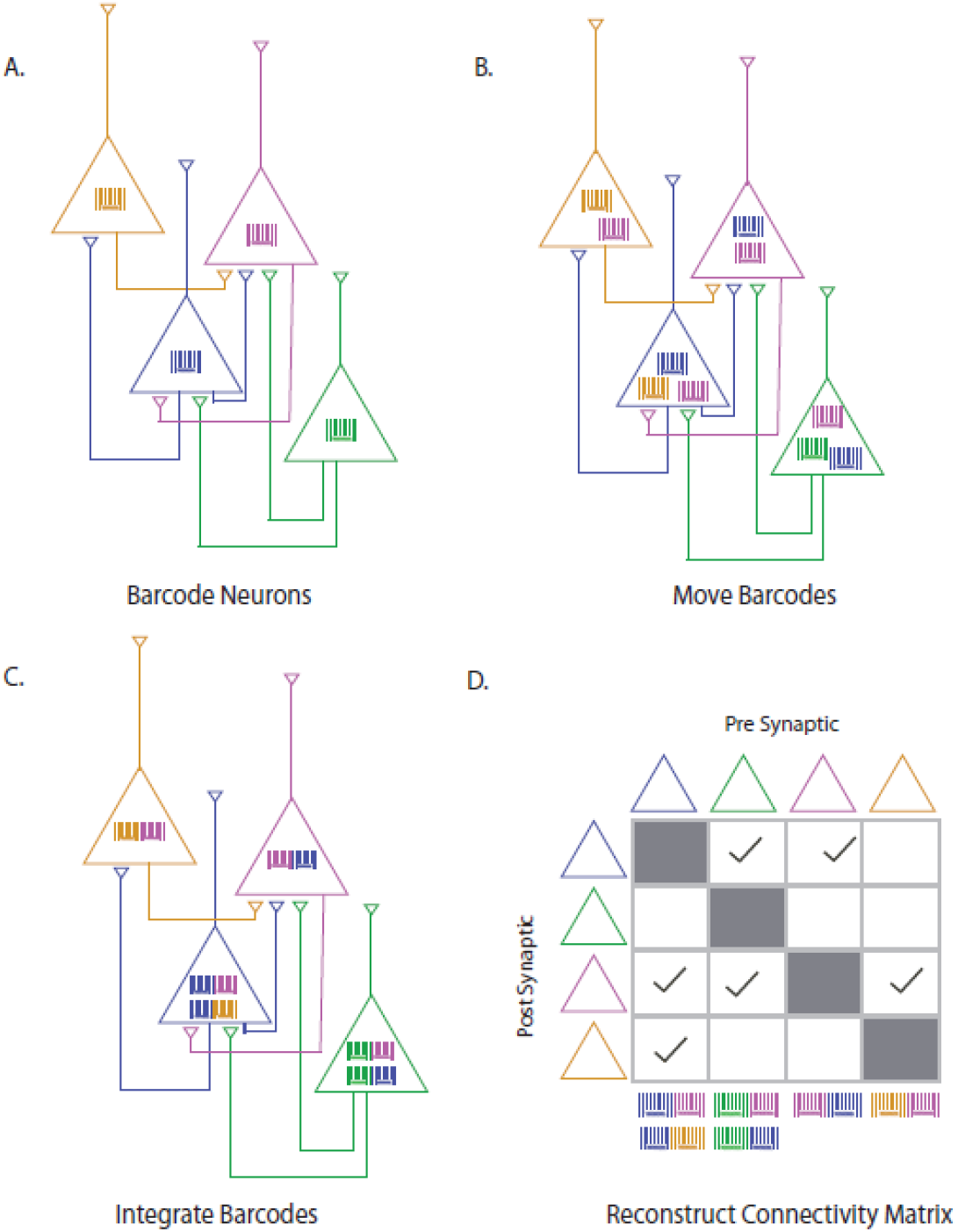
Overview of the approach. (**A**) Individual neurons are marked with DNA barcodes—randomly generated unique sequences of DNA. (**B**) Barcodes move across synapses to connected neurons. (**C**) Barcodes are joined together *in vivo* and then sequenced. (**D**) Joined barcode pairs indicate synaptic partners and can be used to reconstruct the circuit connectivity.

### Barcoding

We first developed a method to barcode each neuron with a random sequence of nucleotides. We used standard shotgun cloning techniques to barcode plasmids *in vitro* (see Methods). Briefly, a synthetic oligonucleotide encoding a sequence of 25 random nucleotides (“N”) was ligated into a PRV amplicon plasmid. The resulting plasmid was then transformed into bacteria and amplified. In practice, this transformation step represented the bottleneck limiting the diversity of the plasmids, with typical diversities in the range of 10^6^–10^7^ (*see Methods*). Barcoded plasmids were transfected into neurons, thereby endowing each transfected neuron with one or more unique barcodes.

### PRV-mediated barcode transport

We next developed a method based on PRV for moving barcodes between synaptically connected neurons. The PRV genome is approximately 150 kb and encodes about 70 genes (Szpara et al., 2011). The replication of PRV begins at the origin of replication (ORI). The PRV ORI_s_ is encoded within a 1.9kb fragment of the PRV genome, previously identified as sufficient to enable helper virus mediated replication of a plasmid (Fuchs et al., 2000; Prieto et al., 2002). The packaging of PRV into the viral capsid requires a PAC sequence (1.2 kb). Under normal conditions the ORI and PAC signal the replication and packaging of the full-length virus, but the ORI and PAC can also signal the replication and packaging of other DNA sequences (Prieto et al., 2002). For example, amplicons containing the ORI and PAC of the closely related herpes simplex virus (HSV), but no other viral sequences, can be grown in high titer in the presence of HSV helper viruses and have been used widely for gene delivery (Fraefel, 2007).

To move barcodes across synapses we engineered PRV amplicons (Prieto et al., 2002) that contain the PRV ORI and PAC sequences. We expected that a PRV amplicon with ORI and PAC sequences transfected into a neuron would, in the presence of a helper PRV, be replicated and packaged and thus be available for movement. Because PRV amplicons are plasmids, they are amenable to the method described above to produce highly diverse populations of barcoded PRV amplicons using standard techniques. Achieving similar barcode diversity in the full-length viral genome, using BAC recombineering or other methods, would have required overcoming significant additional challenges. Although HSV amplicons are widely used to infect neuronal and non-neuronal cells, less is known about PRV amplicons. In particular, the PRV-mediated movement of amplicons across synapses has not been reported.

We first tested the ability of PRV to mediate the movement of PRV amplicons across synapses *in vivo.* We used *in utero* electroporation (E16) to introduce amplicons expressing GFP into layer 2/3 of the left mouse cortex. At day P23 PRV (Becker strain) was injected into the left auditory cortex. After 24 hours neurons labeled with GFP were observed in neurons in diverse brain regions (Fig. 2c). These experiments showed that PRV could mediate movement of amplicons in the brain.

**Figure 2.**
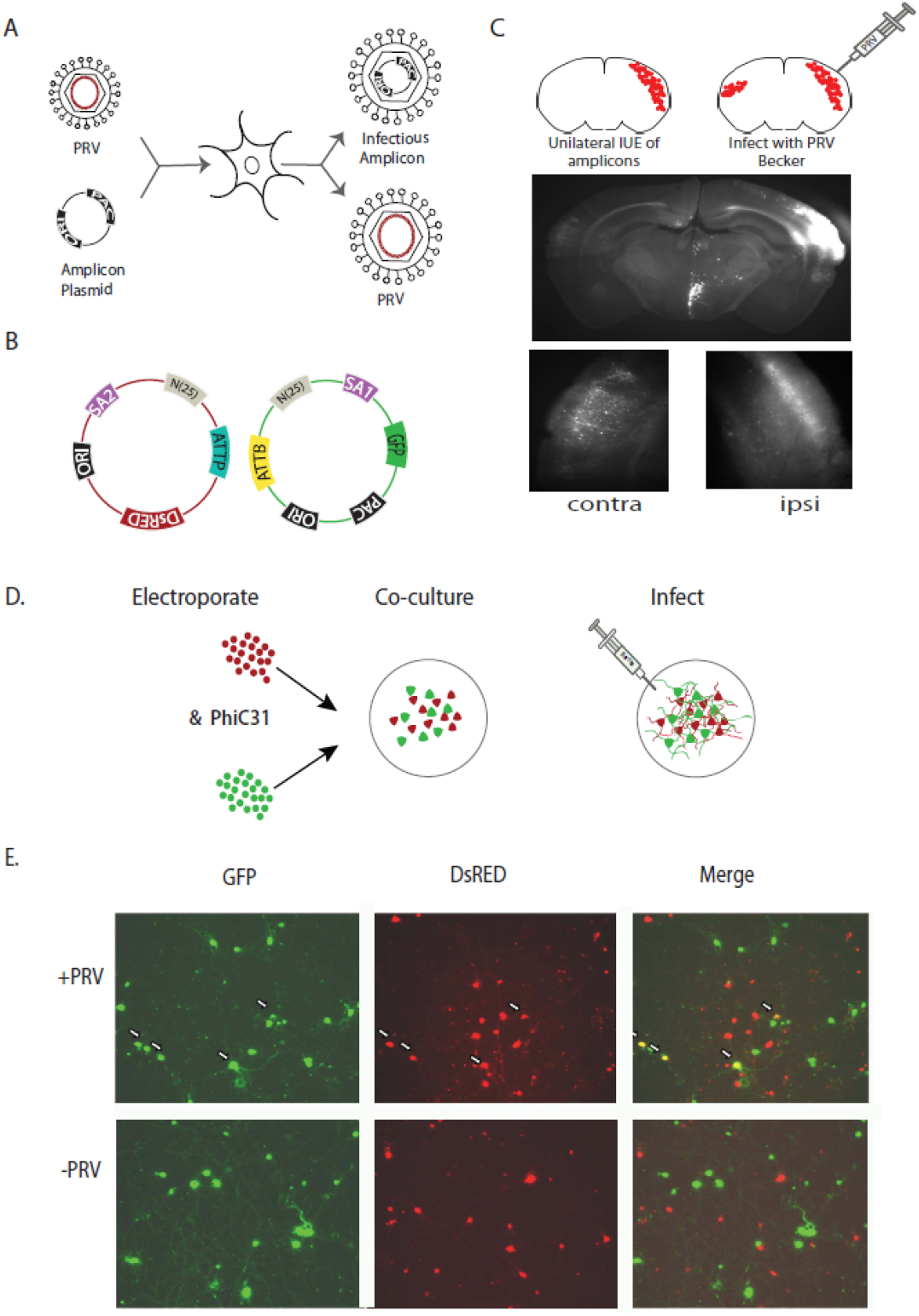
Amplicons undergo PRV-mediated transneuronal movement. (**A**) Viral amplicons are produced when plasmids containing the packaging (PAC) and origin of replication (ORI) sequences of PRV are introduced into the viral packaging cell line in combination with PRV. PRV acts as a helper virus, producing proteins that replicate and package plasmids with PAC and ORI into infectious particles. The output yield after helpervirus infection is a mixture of infectious particles, some containing the viral genome and some containing the amplicon. (**B**) Two types of amplicons were constructed for this study. The *host* amplicon encodes dsRed, PRV ORI and a barcode cassette which includes an Illumina paired end primer binding site (SA2), a randomly generated barcode (N_25_) and a phiC31 integration site (attP). The *invader* includes a GFP, PRV PAC and ORI in addition to a complementary barcode cassette with an Illumina paired end binding site (SA1), a randomly generated barcode (N_25_) and a phiC31 integration site (attB). The absence of a PAC in the host prevents the host amplicons from being packaged and from spreading to connected neurons. (**C**) PRV mediates spread of amplicons *in vivo.* Amplicons encoding dsRed were unilaterally introduced into the right hemisphere of E16 mice using *in utero* electroporation (IUE). At P23 mice were injected with PRV Becker in the right hemisphere causing the replication and spread of amplicons, as evidenced by the presence of amplicons in the contralateral cortex. (**D**) PRV mediates spread of amplicons *in vitro.* E18 cortical neurons were electroporated with either invader amplicon (red) or host amplicon (green) with or without phiC31 and co-plated. PRV Bartha was added at day 3 or 13 after co-plating. (**E**) Images were taken at 24 hours after infection. Overlap of invader and host amplicons was observed in neurons infected by PRV Bartha (upper panel), but not in uninfected neurons (lower panel).

We next tested the ability of PRV to mediate movement of PRV amplicons *in vitro.* We co-cultured two populations of embryonic cortical neurons which had been transfected with two species of PRV amplicons. The first, “host,” species encoded dsRed, and contained an ORI but no PAC sequence, allowing replication but not packaging in the presence of a helper PRV. The second, “invader,” species encoded GFP, and contained both an ORI and PAC, allowing both replication and packaging in the presence of a helper PRV.

In 13-day cultures, many neurons expressed GFP or dsRed robustly, but as expected no neuron expressed both fluorophores, confirming that PRV amplicons require coinfection with helper PRV for movement (Fig. 2e). After twenty-four hours of incubation with the Bartha strain of PRV, co-labeling was observed (Fig. 2e), indicating movement of amplicons. No co-labeling was observed when PRV was added 4 days after plating, before the formation of synapses in these cultures (Weiss et al., 1986), consistent with a synapse-dependent mechanism for PRV spread (Card et al., 1993; Kim et al., 1999). These experiments indicate that PRV can mediate the movement of amplicons *in vivo* and *in vitro.*

### Joining of barcodes

We use phiC31 integrase (Groth et al., 2000) to join host and invader barcodes. PhiC31 integrase is a site specific phage serine recombinase phiC31 that mediates the recombination between the asymmetric attB and attP sequences to form an attL and an attR sequence. Because phiC31 integrase does not mediate the recombination between attL and attR, phiC31 integrase mediated recombination is irreversible, unlike the tyrosine recombinases Cre and Flp.

We constructed invader amplicons in which diverse barcodes were placed near an attB site, and host amplicons in which diverse barcodes were placed near an attP site. Joining of the invader attB site with the host attP site by phiC31 integrase yields a product in which the two barcodes are fused, separated only by an attL site. This joined product can then be amplified by PCR and sequenced (Fig. 3b).

**Figure 3.**
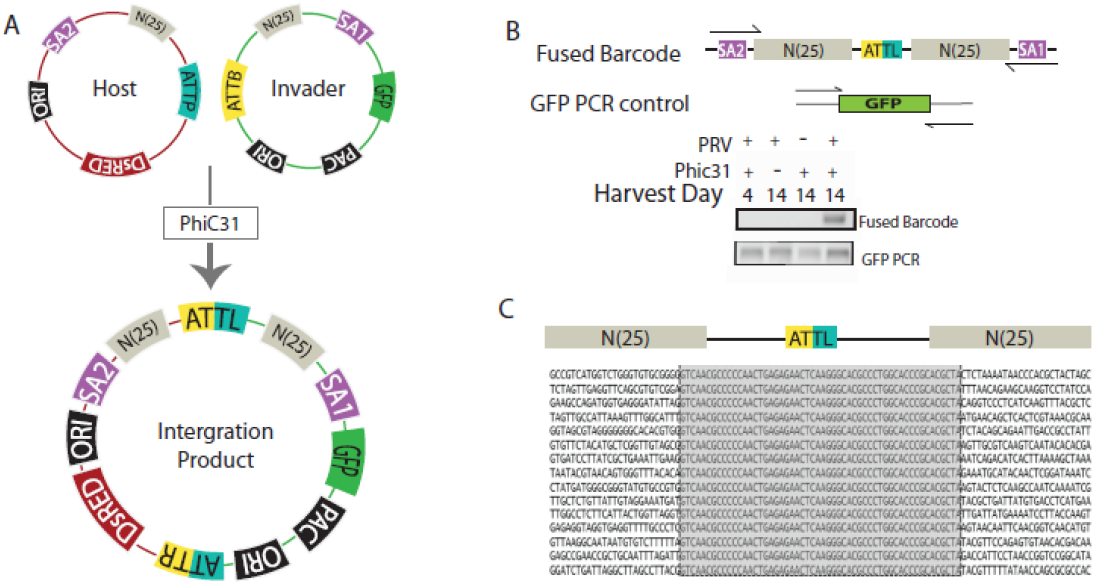
Readout of barcode connectivity by PCR amplification and sequencing of fused barcodes (**A**) Host and donor amplicons are fused by phiC31-mediated integration of the attB on the invader plasmid to the attP on the host plasmid to form an integration product in which the two barcodes are joined and flanked by the Illumina sequencing adaptors SA1 and SA2. (**B**) PCR amplification of the joined barcode (upper panel) and GFP as a control (lower panel). (**C**) DNA sequence of joined invader and host barcodes, with attL sequence highlighted in open box.

To test the ability of phiC31 integrase to join barcodes from synaptically coupled neurons, we adopted a strategy similar to that used above to assess movement of amplicons. However, instead of using the co-expression of different fluorophores, we used PCR amplification of joined barcodes as the readout. Embryonic cortical neurons were transfected with either host or invader amplicons and co-cultured. Host amplicons contained a phiC31 attB site adjacent to a random 25 bp barcode, as well as an ORI for replication. Invader amplicons contained a complementary phiC31 attP site adjacent to a random 25 bp barcode, and both an ORI and a PAC. In some experiments phiC31 integrase was co-transfected.

Twenty-four hours after incubating with PRV, we extracted DNA and amplified the joined barcode product by PCR. Fig. 3a shows that the product of the expected size (223 bp, which includes a pair of 25 bp barcodes, a pair of 4 bp tags, a 58bp SA1 and 61bp SA2 illumina sequencing adaptor and a 46 bp attL site) can be detected, indicating both movement of the invader amplicon and joining of the host and invader amplicon. As expected, no product was detected in the absence of either PRV infection or phiC31 integrase, confirming the dependence of the product on both movement and phiC31-mediated joining. In addition, no product was observed in 4-day cultures, before the formation of synapses in these cultures (Chang and Reynolds, 2006; de Lima et al., 1997), consistent with a synapse-dependent mechanism for PRV spread(Card et al., 1993; Kim et al., 1999).

Conventional Sanger sequencing of 20 colonies of the topo-cloned PCR product confirmed that the joined product had the expected structure, consisting of the invader barcode, the attL site, and the host barcode (Fig. 3c). We therefore performed high-throughput paired-end (PE) sequencing with an Illumina MiSeq.

### Computational algorithms for inferring connectivity

We have implemented a computational pipeline that is capable to efficiently convert sequencing data into connection matrix (Figure 4). The input of this pipeline is the set of sequences of barcode host-invader pairs and attL sites. Here we will call such sequences ‘reads’. High-throughput sequencing yielded approximately 8.3 million such sequences. We applied a series of preprocessing filters to these sequences that implemented several steps of error correction of the barcode pairs. First, we discarded any reads that contained unassigned nucleotides (“N”) at any position. Second, we removed the reads that failed to match the phiC31 attL consensus sequence perfectly. Because high-throughput sequencing is preceded by barcode amplification using PCR, each barcode pair is represented in several copies. As the next step, we identified unique barcode sequences within the list of reads. This operation yielded 376 thousand unique barcode pairs. In our dataset, each unique barcode pair was accompanied by the corresponding number of copies. Many of these reads appeared only in low copy number, consistent with errors in PRV replication, PCR amplification and sequencing. As the third step of error correction, therefore, we considered only reads that occurred in at least 5 copies. This yielded 16,622 unique barcode pairs.

**Figure 4.**
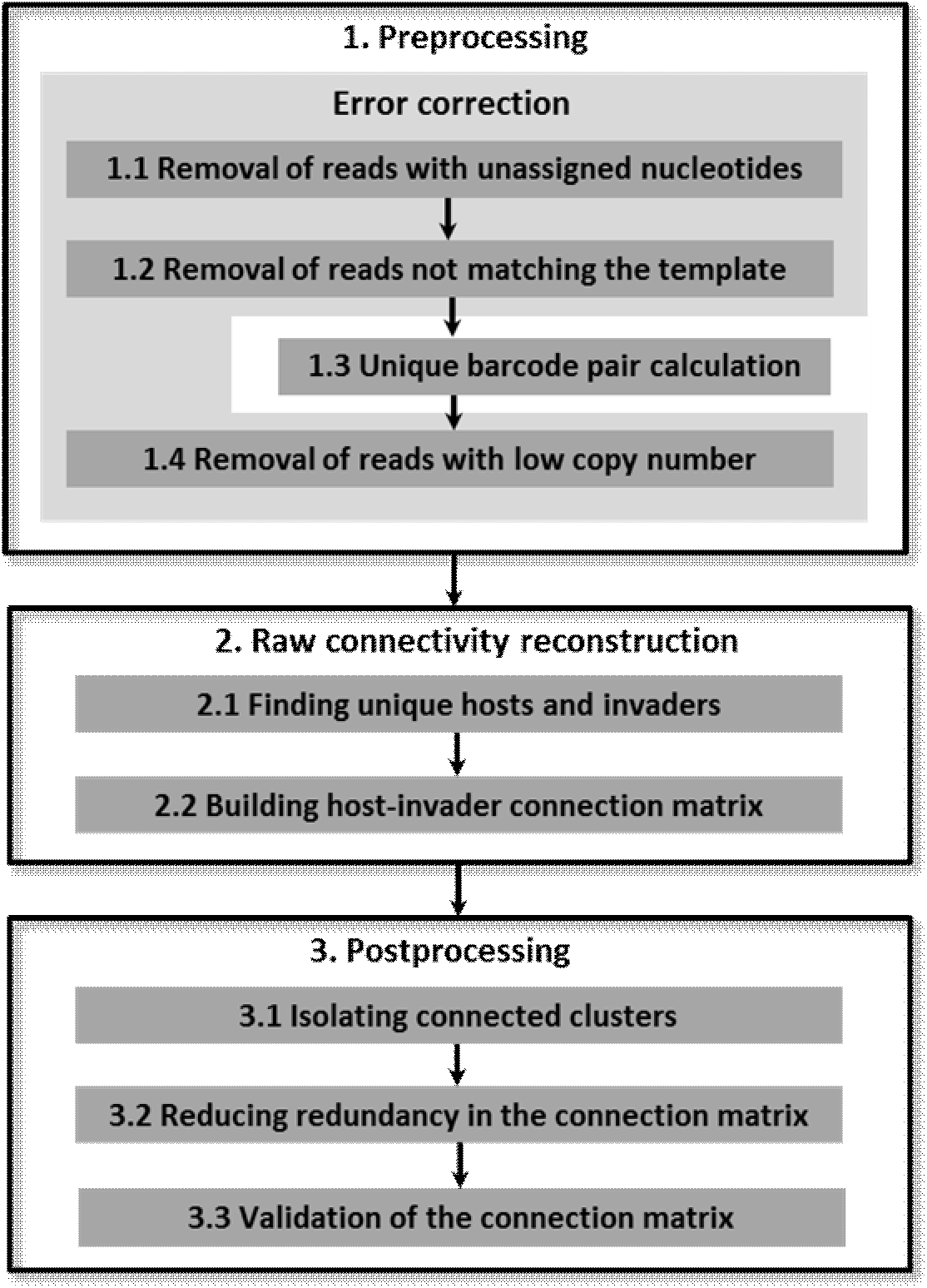
The overview of the computational pipeline for circuit reconstruction.

The next step of our algorithm resulted in the construction of raw connectivity between hosts and invader barcodes. First, we identified unique invaders (n=6,635) and unique hosts (n=9,042) within the list of 16,622 unique barcode pairs obtained in the previous step. Second, we defined a sparse 6,635 x 9,042 raw connection matrix with 16,622 nonzero elements representing the connections from 9,042 hosts to 6,635 invaders. This operation resulted in the connectivity matrix between host and invader barcodes that may reflect real connectivity between neurons (Figure 5A).

**Figure 5.**
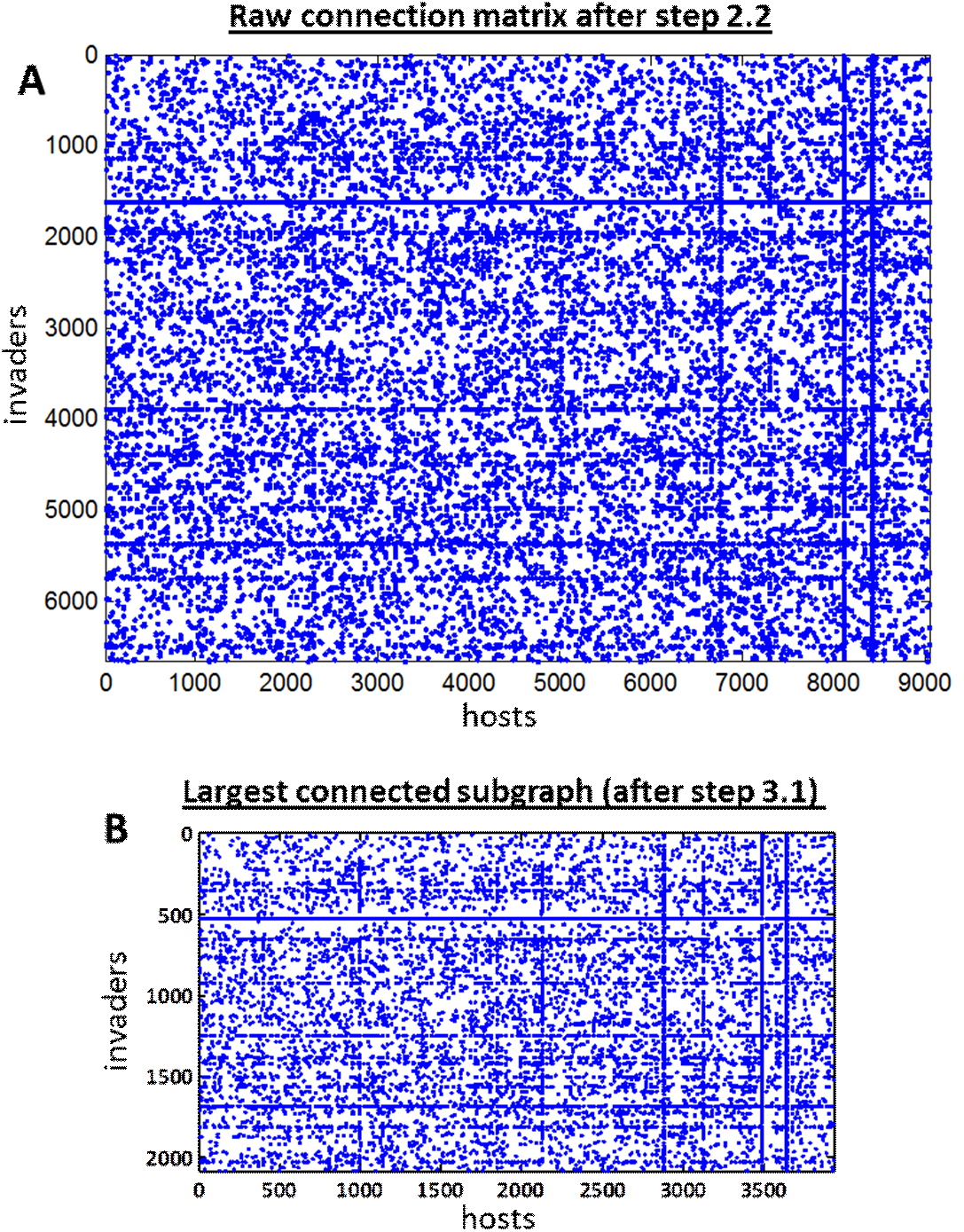
Connection matrices generated by the algorithm. (A) 6,635 by 9,042 raw connection matrix resulting from step 2.2 (Figure 4). Each dot represents a unique invader-host pair. (B) The forest fire algorithm identifies connected subgraphs within the raw connection matrix (step 3.1). Connectivity in the largest subgraph is shown (2,083 by 3,929).

Several additional steps are needed to make raw connectivity understandable as neuronal connectivity. These steps are included in the set of algorithms called postprocessing (Figure 3). First, we identified connected subgraphs within the graph defined by the raw connectivity (Figure 5A) using the forest fire algorithm (Hopcroft and Tarjan, 1971). We found that 16622 connections were organized into 4,077 discrete subnetworks ranging in size from 6,012 to 2 nodes (including both hosts and invaders). The largest connected subgraph contained 2,083 invaders and 3,929 hosts (6,012 nodes in total). The next largest connected subgraph contained 28 invaders and 2 hosts. We limited the subsequent analysis to the largest connected subgraph. The connectivity of this connected subgraph is defined by the matrix *C_ij_*, where the invader index *i* runs between 1 and 2,083, and the host index *j* varies between 1 and 3,929 (Figure 5B).

## Detection of barcoded nodes with correlated connectivity

Within the largest connected subgraph of the raw matrix (Figure 5B), many barcodes had very similar connectivity. This likely resulted from the transfection of multiple invader or host barcodes, with distinct sequences, into single neurons. If all subsequent processes—including PRV-mediated transsynaptic transmission, phiC31-mediated joining, PCR amplification and sequencing—were perfectly efficient and error-free, then multiple barcodes transfected into a given neuron would have identical connectivity. We therefore developed a novel clustering algorithm to identify nodes with similar connectivity, reasoning that high similarity was more likely to occur due to multiple barcoding of a single neuron than to highly shared patterns of connections within the network of cultured neurons. This algorithm is represented by step 3.2 (Figure 4). The details of the clustering algorithm are described below (Figure 6).

**Figure 6.**
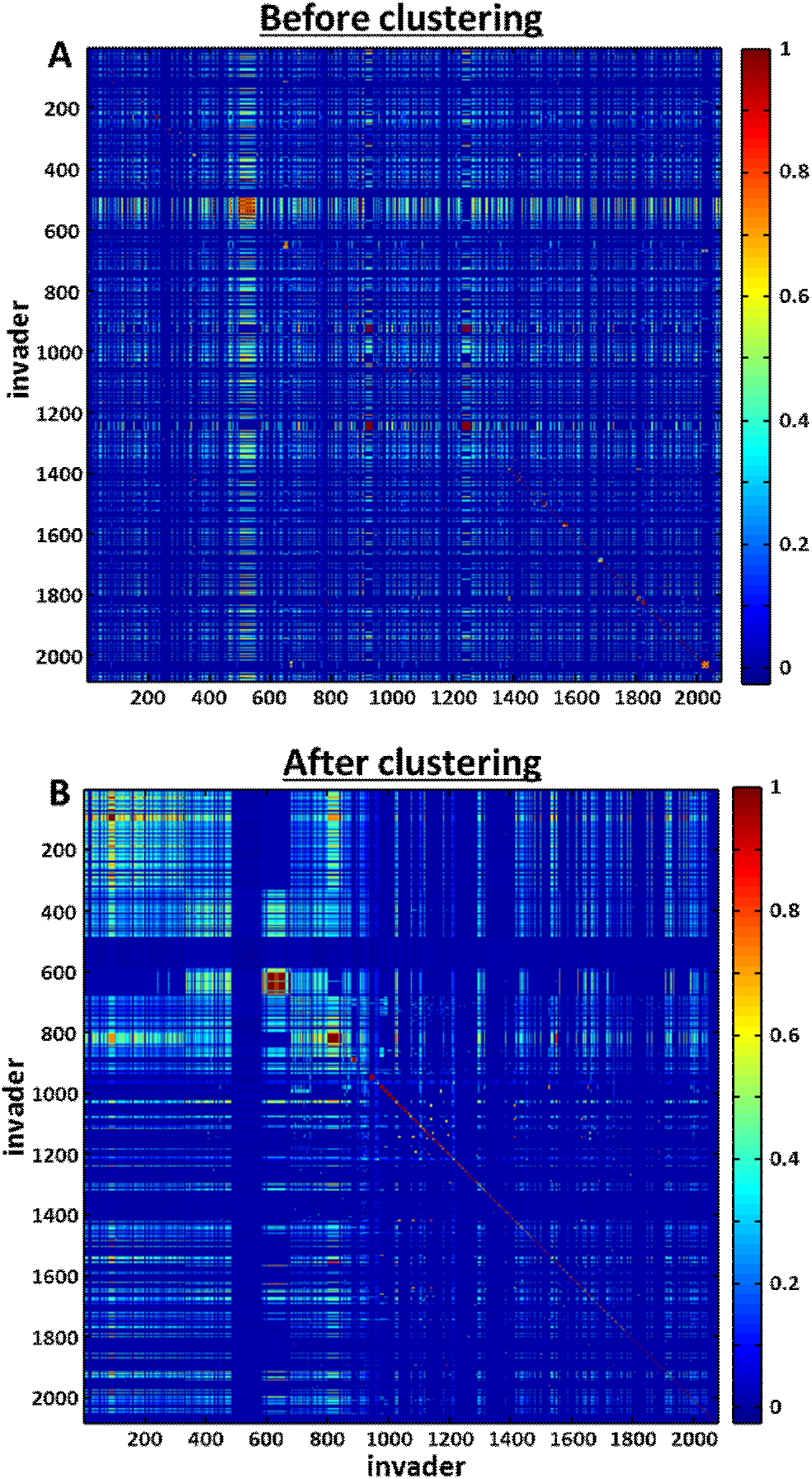
Pearson correlations in connectivity for pairs of invaders. (A) Before clustering. (B) After clustering. The ordering of invader barcodes is changed in (B) compared to (A) so that nearby barcodes belong to the same cluster, i.e. share high similarity in their connectivity patterns to the host barcodes. Clustering is produced by the greedy multidimensional clustering algorithm as described in the text.

To detect neurons with similar connectivity, we calculated the pairwise similarity matrices of connectivity for both invaders and hosts 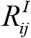 and 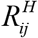. Here 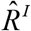 and 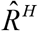 are square matrices, 2,083 x 2,083 and 3,929 x 3,929 respectively. An element of 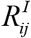 contains the Pearson correlation (the R-value) between connectivity of invaders *i* and *j* (Figure 6A). Similarly, an element of 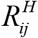 contains Pearson correlation between connectivity of host barcodes *i* and *j*. To evaluate these correlation matrices we used the binary form of invader-host connectivity that is equal to 0 or 1 depending on whether a single barcode pair with this invader and host is present in the sequencing results. To determine which host and invader barcodes belong to the same network node, we used a greedy multidimensional clustering algorithm inspired by watershed algorithm used in morphological image analysis (Beucher and Lantuéj, 1979; Roerdink and Meijster, 2000) and Kruskal algorithm for finding the shortest spanning tree of a graph (Leiserson et al., 2009). We therefore called our algorithm Kruskashed. We describe the application of the algorithm to 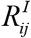 (Figure 7); the application to 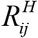 is analogous. First, we identified pairs of invader barcodes that are correlated at the high level of 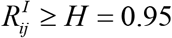 and found a set of connected clusters amongst these pairs. This step is needed to alleviate the effects of noise on clustering results and is analogous to the marker control step described for the watershed algorithms. The connected clusters were used as markers that initiate the set of clusters for the subsequent greedy steps. To practically implement this step, we just set all values of connectivity correlations with 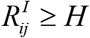 to 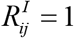 (Figure 7A). Second we grew the clusters by adding pairs of invader barcodes one by one in the order of decreasing correlation 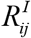. On every step of this iterative procedure, a pair of invaders is considered to be added to the existing clusters. If only one of the invaders within the pair belongs to an existing cluster or if both invaders belong to the same cluster, this pair is attributed to this cluster. If none of the invaders within the pair belongs to an existing cluster, we initiated a new cluster. If two invaders belong to different clusters, the invaders are left untouched. This iterative procedure was repeated until all invaders are attributed to a network node cluster. The same algorithm was used to cluster the host nodes.

**Figure 7.**
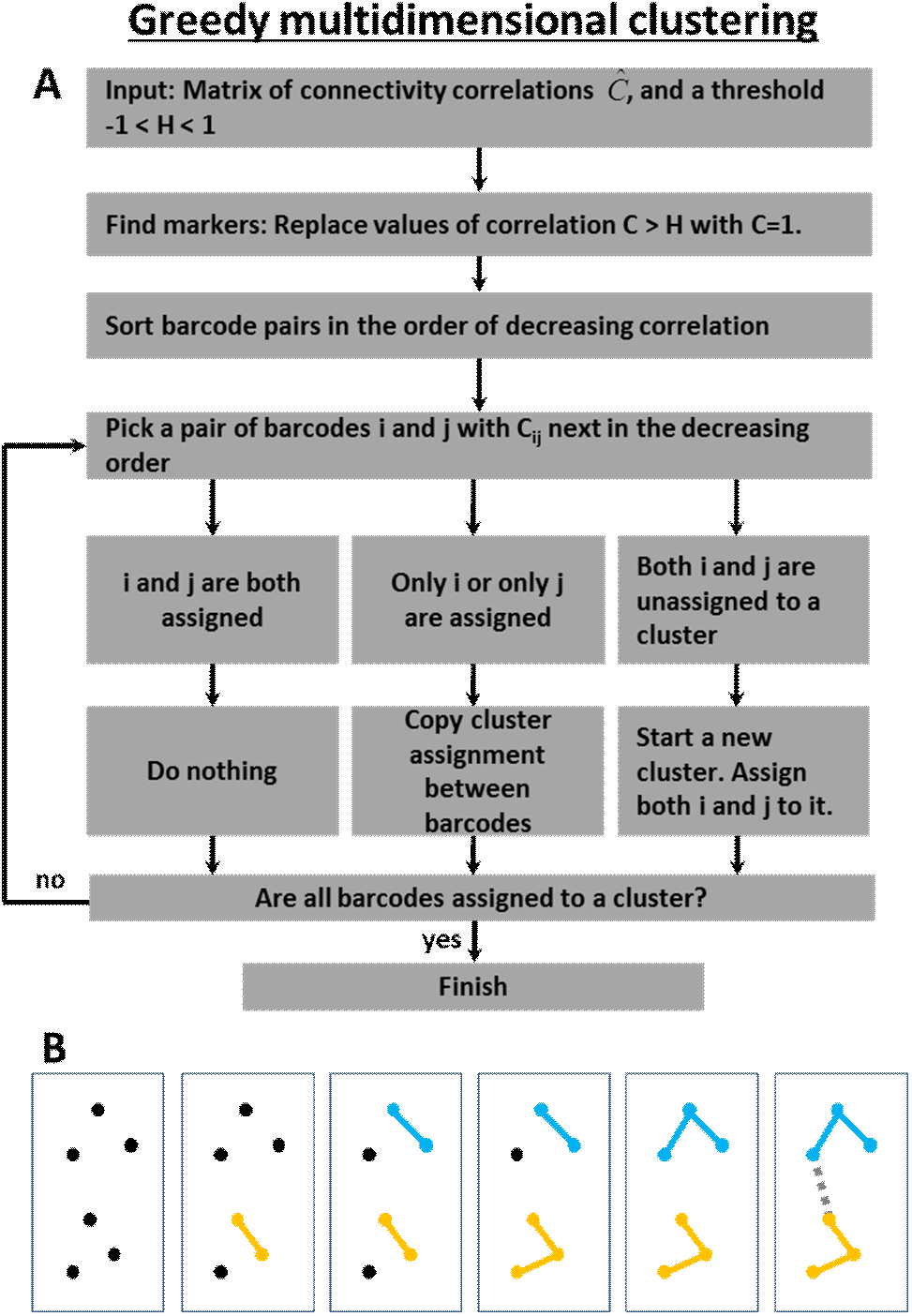
Greedy multidimensional clustering algorithm (Kruskashed) used here to discover clusters of barcodes with correlated connectivity assumed to represent individual neurons. (**A**) The flowchart of the algorithm. (**B**) Illustration of the algorithm clustering a set of six barcodes (dots). The distances between dots represent correlations of connectivity. At each step a pair of barcodes is considered that is next in the order of decreasing distance (correlation) from each other. Three possible scenarios for the pair assignment in (A) are illustrated in the panels two, four, and six.

After applying this redundancy reduction procedure [step 3.2, Figure 4] to the 2,083 by 3,929 invader-host connectivity matrix (Figure 5B), we obtained a connectivity matrix for invader-host network nodes that had dimensions of 456 by 786. We denote this reduced connection matrix *S_ij_*. Indexes *i* and *j* run from 1 to 456 and 786 respectively and represent clusters of invader and host barcodes with similar connectivity. Such connectivity correlations are expected in our network if a cell is electroporated by several host or invader plasmids with different barcode sequences. Although this could not be independently verified, we assumed that these clusters of highly correlated barcodes represent individual neurons that captured several different invader or host barcodes. The reduced connection matrix, *S_ij_*, therefore, attempts to capture connectivity with single neuron precision. The network represented by matrix 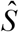 is shown in Figure 8s.

**Figure 8.**
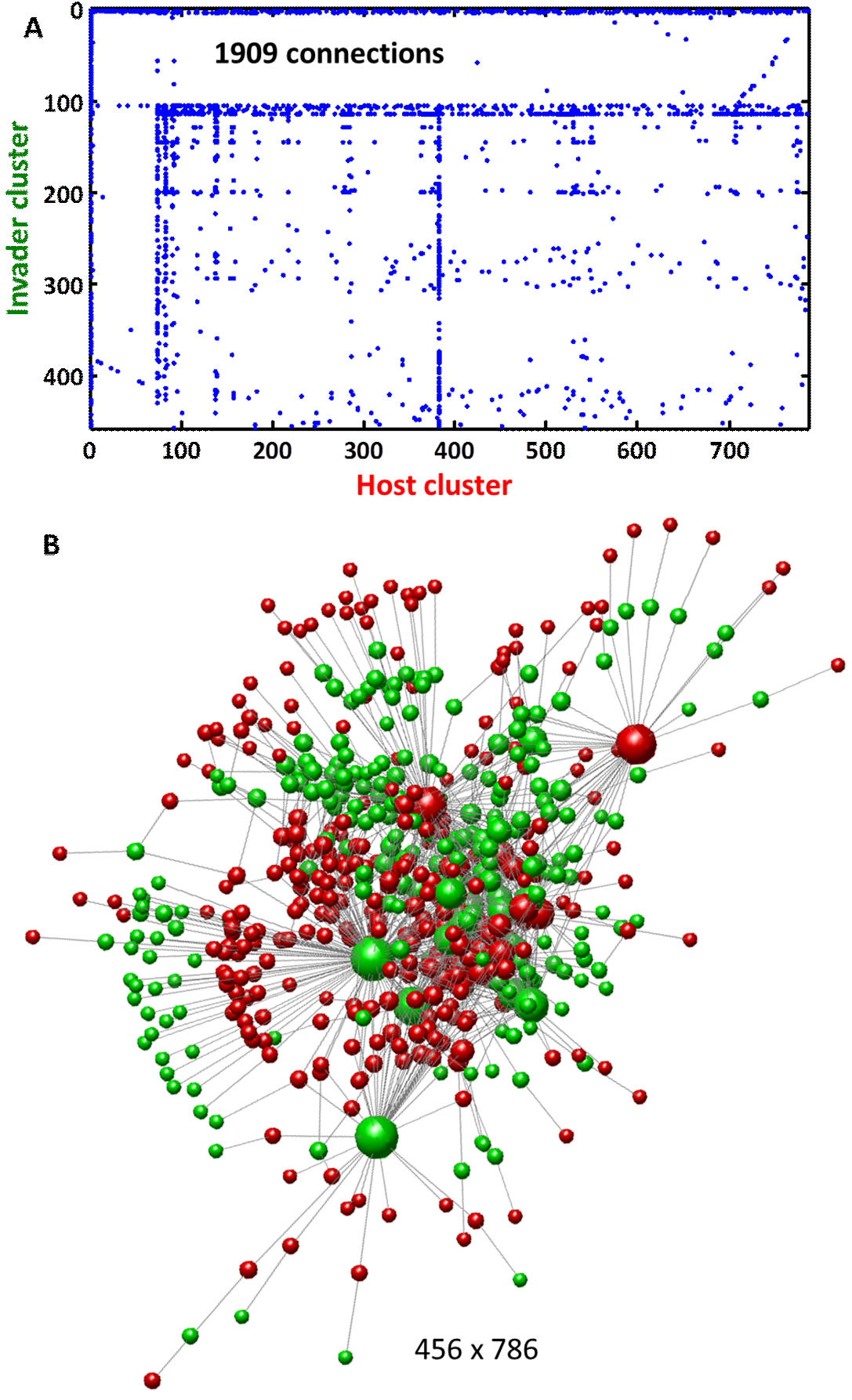
Sequencing results can be interpreted as a network of connected barcodes. (**A**) The connectivity matrix between clusters of correlated barcodes that can be interpreted as individual neurons. (**B**) The graph defined by the connection matrix rendered by the force-directed graph drawing algorithm (green=invaders, red=hosts). The volume of each node represents this node’s coordination number.

### Algorithms for statistical validation of the connectivity matrix

After obtaining connectivity between barcodes we noticed that some invaders had highly similar patterns of connectivity to the hosts, as described above. We have also made a similar observation for the invaders. This phenomenon was interpreted as multiple host or invader barcodes impregnating the same neurons, and, consequently yielding similar connectivity patterns. How similar are the networks rendered from these redundant sets of barcodes? To answer this question, we developed an algorithm that reconstructs connectivity from different non-overlapping sets of barcodes. The purpose of this algorithm is to provide internal cross-validation of individual entries in the connection matrix. If the same entry can be obtained using different sets of barcodes, this connection is likely to exist in reality. It should be noted that the opposite is not true. Our dataset includes many connections that result from a single combination of host-invader barcodes. Such connections are not expected to be replicated in the barcode resampling. Therefore, if a connection cannot be replicated from two non-overlapping sets of barcodes, this connection may still physically exists. Our analysis described here serves to validate but not to invalidate connections.

The details of our algorithm for validation are shown in Figure 9. We resample the sets of invader and host barcodes in two separate steps of the algorithms 500 times each. This is done to avoid recomputing Pearson correlation matrix (Figure 6), the longest step of the algorithm. Within each resampling step, we split one barcode set, say invaders, into two non-overlapping random subsets. We use the full invader-to-host connectivity matrix (Figure 5B) for the subsequent steps. We then repeat the connectivity reconstruction procedure, including obtaining correlation matrix, clustering based on correlation, and computing the cluster-to-cluster connectivity matrix, for both randomly selected non-overlapping subsets of invaders. The simplification that helps streamline the algorithm is that for invader resampling, *host* clusters and invader-invader correlation matrix do not need to be recomputed. We then align invader barcode clusters obtained for both randomly selected sets to the clusters obtained for the full set of barcodes. To this end, for each resampled cluster we find the full set cluster with the higher overlap. This procedure allows us to identify clusters between two randomly selected subsets of barcodes and to compare the resulting connection matrices. After repeating this procedure 500 times for invader and host barcodes we find the fraction of matches, i.e. the fraction of 1000 resamples in which connectivity rendered on the basis of randomly chosen and non-overlapping sets of barcodes is the same. Our procedure cannot generate new connections between existing clusters, therefore, zero connectivity yields no matches. The matrix of matches that we call the Q-matrix is shown in Figure 10.

**Figure 9.**
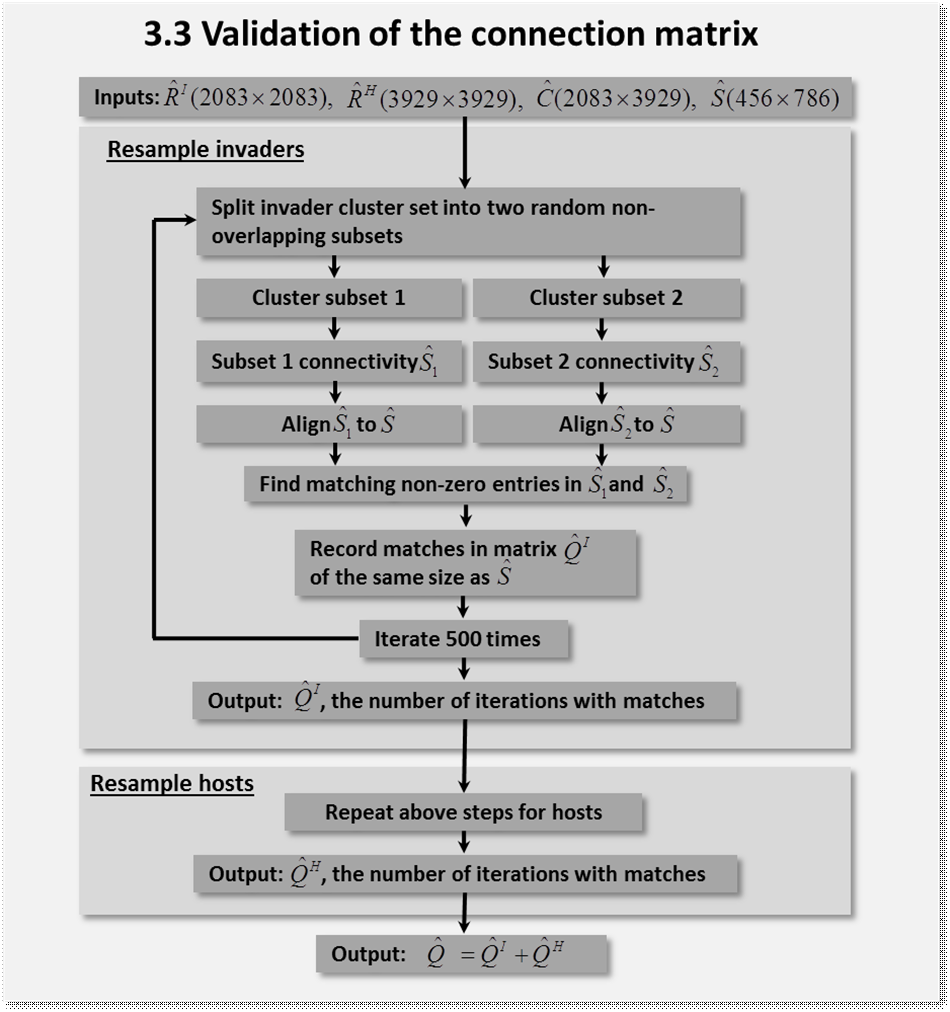
The flowchart of the algorithm used here for internal validation of the data.

**Figure 10.**
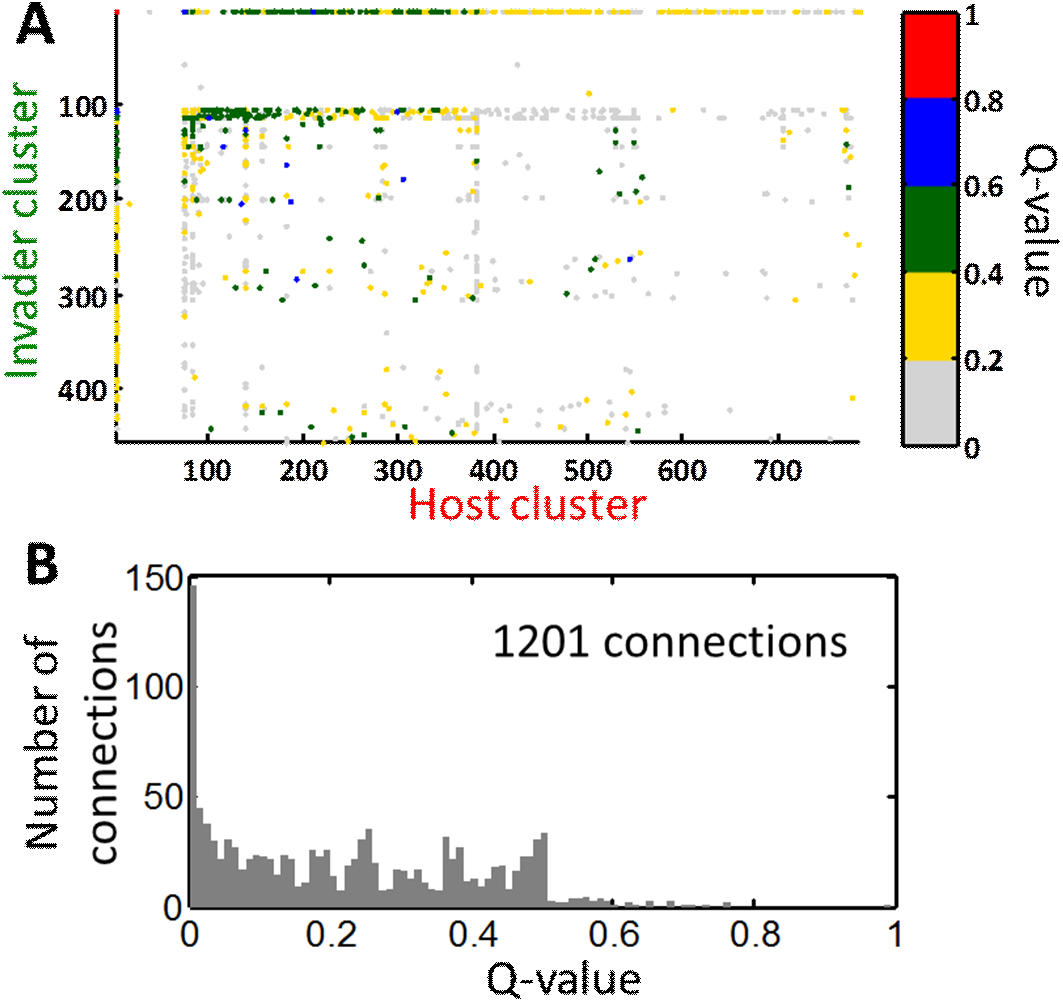
The results of internal validation. (**A**) The number of matches between reconstructed connectivity matrices obtained from split datasets (Q-value). (**B**) the distribution of non-zero Q-values in (**A**).

The values of entries in the Q-matrix that we call the Q-values represent the reproducibility of our network when physically different sets of barcodes are used to reconstruct its structure. High Q-values signify connectivity that is highly reproducible. However, low Q-values do not mean an insignificant connection, because we expect that substantial number of connections is produced by single barcodes electroporated into individual neurons.

## Discussion

Here we have described the first steps of our method BOINC, demonstrating how neural connectivity can be converted into a problem amenable to high-throughput DNA sequencing. We show that (1) DNA barcodes can be introduced into neurons; (2) these barcodes can be moved by PRV to synaptic partners; and (3) barcodes from connected neurons can be joined and sequenced using high-throughput sequencing technologies. We also describe the set of computational algorithms that produces connectivity on the basis of sequencing data. These results lay the groundwork for applying the tremendous potential of DNA sequencing technologies to neural connectivity.

The choice of PRV was motivated by several considerations. PRV has been widely used to trace neuronal circuitry, and so was a good candidate to move genetic barcodes across synapses. Because PRV is a DNA virus, we could exploit phiC31 integrase to join DNA barcodes. Although other viruses including rabies can also move genetic material across synapses (Wickersham et al., 2007), rabies is an RNA virus and thus would have required a means to join RNA barcodes *in vivo* that are not available at this time. Although PRV may not be the ideal candidate for mapping the complete connectome—for example, guaranteeing that every neuron expresses PRV would require generating transgenic animals encoding the viral genome—PRV is ideally suited to mapping long-range connections, which are particularly challenging for microscopy-based approaches.

It is important to note that connectivity reconstructed by the PRV-based method is difficult to interpret as neuronal connectivity with a single synapse precision. The lack of correspondence between barcode-to-barcode and neuronal connectivity may arise from several factors. First, in this experiment we used a replicating PRV strain (Bartha) capable of mediating polysynaptic barcode movement. Some of the invader barcodes may therefore have passed through more than one synapse before joining with an immobile host barcode. Second, we cannot rule out the viral particles passing via non-synaptic contacts between cells. Although, in the imaging assays, we observe no invaders penetrating into host cells when PRV was added 4 days after cell plating, i.e. before synaptic contacts are formed in vitro, it is impossible to fully rule out non-synaptic barcode transmission in the sequencing-based approach. Third, because barcodes were introduced into neurons through transfection, a method that typically delivers many plasmids to each recipient cell, each neuron may be represented by several nodes in the graph. Although we have developed a computational clustering algorithm that discovers such occurrences, we were not able to independently confirm the accuracy of our computational findings. Because these challenges remain at present unresolved within PRV-based method, the network depicted in Figure 8B cannot be interpreted as a neural circuit, but must instead be treated as a formal graph describing the interaction of barcodes. It is these considerations that led us to move to a second generation experimental system, synseq (Peikon et al., 2017).

**Figure 11.**
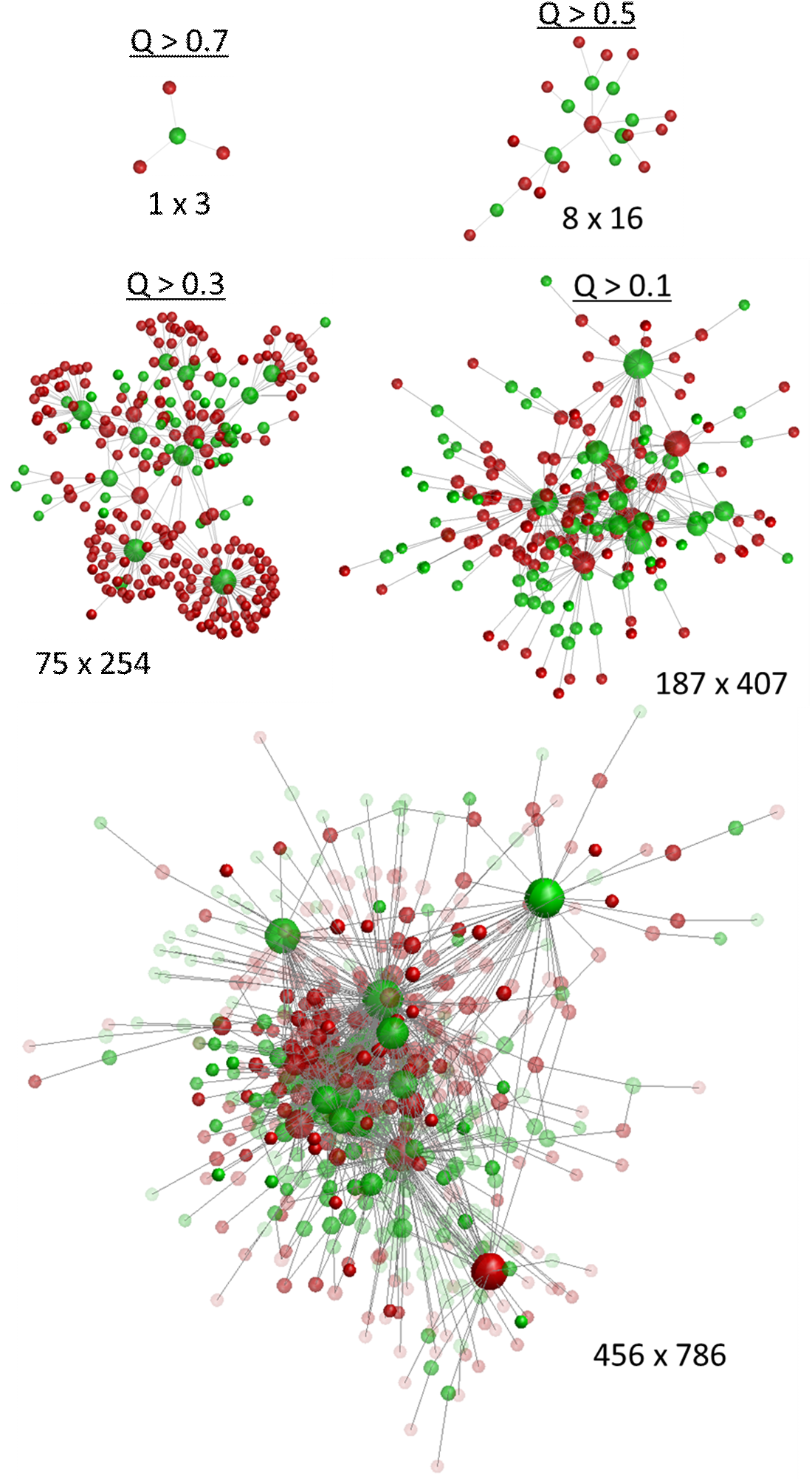
The connectivities rendered by force-directed graph drawing algorithm at different levels of Q-values, as indicated. Bottom: The entire connectivity combined with Q-values indicated by the nodes’ transparency.

An additional important challenge will be to assess the accuracy of the circuits obtained using sequencing methods. All methods have false positives (i.e. inferred connections that do not exist) and false negatives (actual connections that are missed). The ideal solution would be to compare the circuit obtained using sequencing to a “ground truth” circuit obtained using other methods. Although the ground truth circuit is available in one case, *C. elegans,* PRV does not appear to propagate in invertebrates. Thus other approaches will need to be developed to validate this approach.

Despite these limitations, we have demonstrated the feasibility of each of the core components needed to convert neural mapping into a form amenable to DNA sequencing. In addition, we have developed a set of computational algorithms that allows to reconstruct connectivity from sequencing data. Our methods yielded a network that is described by a 456-by-786 connectivity matrix between putative neurons (Figure 8), which is the largest neural network reconstructed so far. This connectivity was obtained at the fraction of the cost and using a fraction of computational effort compared to conventional methods, such as electron microscopy. By encoding the connectivity of neural circuits intro DNA sequence and exploiting high throughput DNA sequencing technology, low cost circuit sequencing could be used as a routine screening method to explore the dynamics of brain circuits in health and disease. This approach could pave the way to uncover the detailed architecture of neural networks and a better understanding of neural circuit function.

